# Stingless Bees of the Amazon Forest: Taxonomic and Geographic Gaps and the Potential for Meliponiculture

**DOI:** 10.1101/2025.07.15.664956

**Authors:** Aline C. Martins, Tais M. de A. Ribeiro, Thais Vasconcelos

## Abstract

Insects, especially those distributed in tropical forests, are greatly impacted by shortfalls in biodiversity knowledge. It is no surprise, then, that there are considerable gaps of knowledge on geographic distribution and taxonomy in the most diverse group of Amazonian bees, stingless bees (tribe Meliponini). Addressing these knowledge gaps is crucial for directing future research efforts and funding effectively, especially given the relevance of meliponiculture in the local economy. Here, we assess the geographic and taxonomic gaps in knowledge for Meliponini in the Amazon basin. We found that there are 27 genera and 239 valid species of native Meliponini in the region, but, among these, 78 species have five or less occurrence points available in open datasets of biodiversity data, while 7 have no available occurrence data. This issue stems from both a lack of collection efforts—especially in areas far from rivers and roads—and insufficient digitization of existing collections. We argue that future research should prioritize resolving taxonomic uncertainties in species complexes that suffer from both geographic and phylogenetic knowledge gaps. Some of these species are also important for meliponiculture, such as *Melipona* (*Michmelia*) *seminigra* and *Tetragonisca angustula*. Besides, to overcome the large geographical shortfall, both digitization and new collection effort should be employed, including new standardized methods to access the canopy, the largest frontier of biodiversity knowledge in the Amazon.

## Introduction

Large-scale knowledge in ecology and evolution is often limited by seven well-defined shortfalls: species taxonomy (Linnean), distribution (Wallacean), abundance (Prestonian), evolutionary patterns (Darwinian), traits (Raunkiæran) and biological interactions (Eltonian) (Hortal et al. 2015). These biodiversity knowledge gaps hinder ecological and evolutionary research, while simultaneously limiting our ability to inform conservation initiatives (Cardoso et al. 2011). Acknowledging and quantifying the extent of these gaps can both enhance our capacity of describing biodiversity and improve our ability to predict how it might change in the future, since resources can be more effectively directed towards where the gaps are (Hortal et al. 2015).

Due to the heterogeneity in species geographic distribution, as well as uneven survey efforts and research infrastructure, the availability of biodiversity data varies widely across both space and time (Hortal et al. 2015). Additionally, some organismal groups are more data deficient than others. Insects, for instance, exemplify major shortfalls, largely due to their incredible diversity of species (Entomological Society of America 2021). In this sense, insect biodiversity in the largest tropical rainforest in the world, the Amazon, not surprisingly, suffers severely from all of the main shortfalls in knowledge. The Amazon basin spans approximately 6.7 million km^2^ and has been the focus of many studies in recent decades, but ecological research in the region remains restricted to easily accessible areas close to roads and rivers. Knowledge gaps persist across more than 50% of the upland (*terra firme*) areas of the Amazon basin (Carvalho et al. 2023), making this region one of the largest frontiers for biodiversity research. This tropical biome has also been a major source of biodiversity in the Neotropics (Antonelli et al. 2018), largely due to the region’s long evolutionary history, shaped by a combination of climatic and geological changes (Hoorn et al. 2010).

Stingless bees (Hymenoptera, Apidae, Meliponini), an important exception to the inverse latitudinal gradient in bee biodiversity (Michener 1979, Orr et al. 2020), is primarily distributed in tropical and subtropical regions of the globe, with a peak in species richness in the Amazon region (Michener, 1979). Of the about 550 species of stingless bees known globally, 77% occur in the Neotropics (Grüter 2020) and most are strongly associated with the Amazon Basin (Melo 2020). They represent the most diverse group of corbiculate bees – those possessing a pollen basket to carry pollen (Martins et al. 2014) – and the most diverse group of eusocial bees (Michener 2007). Given the crucial ecological roles played by insects, and particularly by bees as major pollinators of flowering plants (Ollerton et al 2011; Martins et al., *in review*), it is fundamental to reduce the knowledge gaps about them, beginning with taxonomic studies that better delimit species (Linnean shortfall), systematic studies that address the relationships among them (Darwinian shortfall) and descriptions of their geographic distributions (Wallacean shortfall). Reducing the knowledge gaps in these areas will, in turn, provide a solid foundation in which to address the other shortfalls mentioned earlier, which are significant for bees in general (Marshall et al. 2024).

Stingless bees play a fundamental role in the pollination of tropical forests (Bawa et al. 1985, van Dulmen 2001, Martins et al. in review) and interact with a large diversity of plant families worldwide (Bueno et al. 2023). On flowers, they forage not only for pollen and nectar, but also for resins, oils and waxes (Roubik 2023). Although often considered generalist flower visitors, like many eusocial bee species, they tend to prefer foraging in trees and mass-flowering plant species (Martins et al. 2023). Beyond their ecological significance, stingless bees also have an economic and cultural importance given their role in meliponiculture (i.e. stingless beekeeping) (Jaffé et al. 2015, Quezada-Euan & Alves 2020). Meliponiculture is an ancient practice, especially in the Americas and has gained more attention in the last decades due to its relevance in sustainable development (Jaffé et al. 2015). However, shortfalls in knowledge, especially the Linnean and Wallacean, continue to hinder the advance of meliponiculture in the Amazon region (Carvalho-Zilse & Nunes-Silva 2012). These two shortfalls play fundamental and interconnected roles in understanding how biodiversity is exposed to human-induced threats (Baranzelli et al. 2023).

In this study, we address the geographic distribution and taxonomy of Amazonian stingless bees and identify key areas for future research aimed at reducing Linnean, Darwinian and Wallacean shortfalls. Specifically, we aim to answer: 1. What is the taxonomic diversity of stingless bees in the Amazon? 2. What is the extent of the Linnean, Wallacean and Darwinian shortfall in Amazonian stingless bees and which taxa and areas should be the focus of future research? 3. How are the species used in meliponiculture distributed across the Amazon? Finally, we discuss ongoing challenges for research on Amazonian stingless bees, with a focus on taxonomy, biogeography, and meliponiculture.

## Material and methods

### Study design

Our study followed a multi-step approach that consists of: 1. Compiling all available geographic records from public databases; 2. Cleaning the records to remove errors in the coordinates (e.g. coordinates in the ocean); 3. Checking taxonomy validity status of the recorded species using the Neotropical Bee Catalogue (Camargo et a. 2023), the most comprehensive taxonomic reference for bee fauna in the region; 4. Mapping the occurrence of stingless bees within the Amazonian Forest biome, including its various ecoregions according to the WWF; and 5. Conducting a review of taxonomic or phylogenetic studies on Neotropical stingless bees and compiling a list of species currently managed for meliponiculture.

### Geographic distribution data

To characterize the distribution of Meliponini bees in the Amazon in order to characterize the extent of the Wallacean shortfall in the region, we downloaded data from SpeciesLink (CRIA 2024) and GBIF using the search methodology described as follows. In SpeciesLink, we searched for the family “Apidae”, since this database does not include a specific field for “tribe”, thus precluding a query for “Meliponini”. Next, we filtered the resulting dataset to include only Amazonian records, using the mapBiomas (Souza et al. 2020) tool implemented in SpeciesLink. SpeciesLink is based in Brazil, but to make sure we obtained data from all the Amazonian countries, we downloaded the observations for Bolivia, Colombia, Ecuador, Guyana, French Guiana, Suriname, and Venezuela as well. We obtained 57,374 Meliponini records from SpeciesLink.

We downloaded data from GBIF with the package *rgbif* (Chamberlain et al 2025) implemented in *R vs. 4.3.3* (R Core Team 2024). We performed individual searches for the taxon keys for each Meliponini genera according to the Neotropical Bee Catalogue (Camargo et al 2023) and filtered for only records with georeferenced occurrences in South America, and no geospatial issues (*i.e.* invalid, out of range, or country mismatched coordinates). This resulted in 152,476 observations. Both “preserved specimens” and “human observation” records (e.g. those coming from iNaturalist) were kept in the dataset. We combined these datasets using *Tidyverse* (Wickham et al. 2019) data wrangling package in R. We then cleaned the dataset using the pipeline in the package *CoordinateCleaner* (Zizka et al 2019). We used the standard flags in the package, removing coordinates that referred to country capitals (3,890 records), country centroids (236 records) and at sea (3,728 records). Then we removed observations not identified to species. After also removing duplicates, we had 123,249 records, from which 14,383 are unique coordinates per species.

We then filtered the dataset for only observations occurring in the Amazon with the package *sf* (Pebesma & Bivand 2023, Pebesma 2018) and the Ecoregions 2017 shapefile of the World Wildlife Fund (WWF) ecoregions (Olson et al 2001, Dinerstein et al 2017). We selected ecoregions in the amazon based on the Global 200 ecoregions (Olson & Dinerstein 2002). We first selected the biome Tropical & Subtropical Moist Broadleaf Forests occurring in South America and removed the ecosystems related to the Atlantic Forest and Andean forests (Central Andean Yungas, Coastal Venezuela Montane Forests, Chocó-Darién Moist Forests, Talamancan-Isthmian Pacific Forests, Northern Andean Montane Forests). We also included the Savannas occurring in the Amazon Forest Domain (Guianan Savanna and Beni Savanna) to account for the diversity of vegetation in the region.

### Corrections on taxonomic accuracy

To review and confirm the validity of taxonomic information (i.e. species names) compiled in our dataset, we cross-checked this species list names with those available through the Neotropical bee catalogue (Camargo et al. 2023), considered the most trustworthy and up to date source of taxonomic information for Neotropical bees. Species names were checked for correct spelling and validity status at the same time, also considering subspecies as in the case of some *Melipona* species. Those species names retrieved in our data compiling were then classified as 1. valid; 2. valid but misspelled 3. synonyms and 4. *nomen nuda*, i.e. refer to names never formally described in the taxonomic literature.

### Review of taxonomic and phylogenetic studies

To address the Linnean and Darwinian shortfalls for Meliponini bees in the Amazon altogether, we conducted an extensive literature search in June 2024 to assemble a database of taxonomic and/or phylogenetic studies with Meliponini genera occurring in the Amazon. We searched the Web of Science, Google Scholar and Scielo platforms to retrieve publications using the keywords: “phylogeny” OR “taxonomy” associated with taxon names, i.e. “Meliponini” OR the names of the Meliponini genera according to Camargo et al. (2023), for example “Melipona” OR “Trigona”. Each accepted genus name with occurrence in the Neotropics was assessed in this search. To ensure a completeness of our search, we also searched for backward citations, which means we have checked all the referenced papers in each study to assure we were including historical papers, not always retrieved on the database searches. All taxonomic or phylogenetic studies were compiled disregarding specific methods (e.g. if molecular or morphological data were used as the main source of data for phylogenetic inference). Our focus lay on complete taxonomic revisions of genera or species groups covering all species described in each genera.. We decided not to include isolated species descriptions, since our focus laid on pointing out lineages that have complete taxonomic assessments. Potential new species were compiled from Pedro (2014), which based her interpretation on the Neotropical Bee Catalogue and specimens housed in the RPSP – Coleção Entomológica Prof. J.M.F. Camargo, FFCLRP/USP.

### Compiling a list of species used in the meliponiculture

We compiled all species in the National Catalogue of Stingless bees from Brazil (Brasil, 2021), a governmental initiative that pulled together a list of species used for meliponiculture in the country, including their native geographical distribution. As Brazil holds 60 % of the Amazon (IBGE 2019), this list is a comprehensive picture of the species used for meliponiculture in the Region and offers the bases of regulatory practices and management recommendations in the Amazon territory (Menezes et al. (2023). The species present in this catalogue are henceforward cited as “managed” species (in contrast to “non-managed” species, those not used for meliponiculture). We then map the distribution of these species in the region using our filtered dataset of occurrence points.

## Results

### Diversity of managed and non-managed species of stingless bees in the Amazon

We recorded 232 valid species names in 27 genera present in the geographic databases in the Amazon region (Table S1), plus 71 species names found to present underlying taxonomic issues. Specifically, 45 species names were classified as *nomina nuda*, primarily associated with *Scaptotrigona* (15), *Trigonisca* (11), *Frieseomelitta* (5), and *Plebeia* (5). Fifteen additional names were considered synonyms of valid species, and 11 names were identified as misspellings.

*Melipona* is the most speciose genus with 41 valid species and subspecies names, followed by *Trigona* (26), *Partamona* (23), and *Paratrigona* (20) (Figure 1). The remaining genera are represented by fewer than 20 species (Table S1).

**Figure 1.**
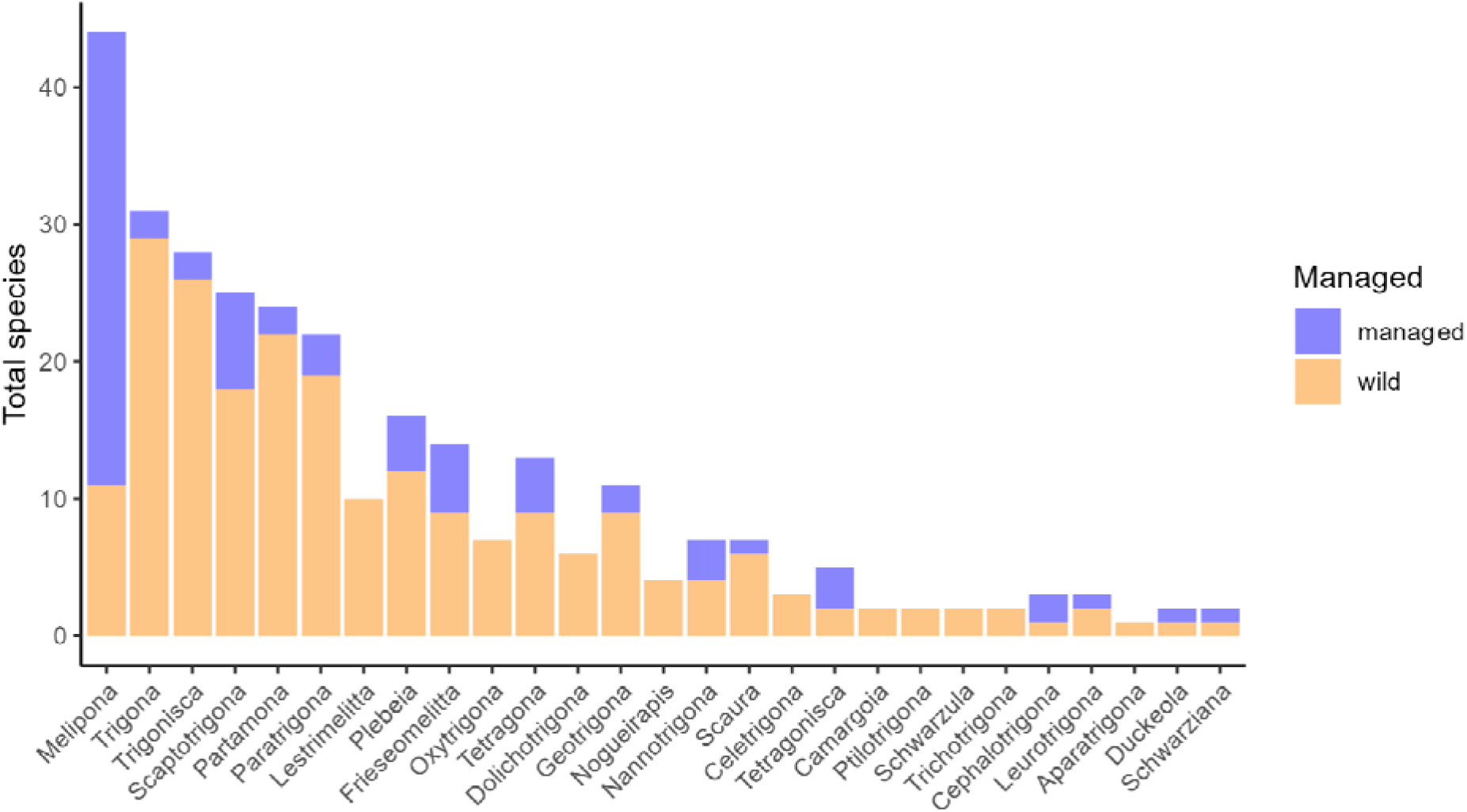
Species of Meliponini per genus in the Amazon organized from the most speciose genus (Melipona) to the least speciose (Schwarziana). Colors indicate the number of managed and non-managed (wild) species for meliponiculture in the Amazon Region.

Among the managed species, more than 50% have records for the Amazon forest (Table S2). *Melipona* is the most prominent genus with about 33 species used in meliponiculture, representing about 78% of the total richness of this genus in the Amazon. Despite being species-rich, *Trigona* is represented in the meliponiculture by only two species, and *Partamona* has no species currently used (Table S2). Other genera with several species used in meliponiculture include *Scaptotrigona* (7 species), *Frieseomelitta* (5 species), *Plebeia* and *Tetragona* (4 species each). Genera such as *Paratrigona*, *Tetragonisca*, and *Nannotrigona* also show moderate representation, with 3 species used. In contrast, genera like *Scaura*, *Leurotrigona*, *Duckeola*, and *Schwarziana* each have only 1–2 species used in meliponiculture. Several genera—*Aparatrigona*, *Camargoia*, *Schwarzula*, and *Ptilotrigona*—have no known representatives used in meliponiculture.

### Linnean and Darwinian shortfall in Amazonian stingless bees

In total, we retrieved 16 studies on taxonomic reviews and/or phylogenetic analysis of complete genera, which are summarized in Table S3. Among the Meliponini, 15 out of the 38 neotropical genera and subgenera (in the case of *Melipona*) have been the focus of taxonomic revisions and/or phylogenetic studies (Table S3), and all of these taxa occur in the Amazon. However, the Linnean shortfall remains substantial: 23 genera or subgenera still lack any complete published taxonomic treatment. Notably, some of the most diverse lineages—such as *Trigona* and *Trigonisca*—exhibit the most pronounced Linnaean shortfalls, with potential new species yet to be formally described (Table 1). Some genera, such as *Plebeia* and *Scaptotrigona*, are currently under taxonomic revision, although these works remain unpublished formally (Table 1).

**Table 1.**
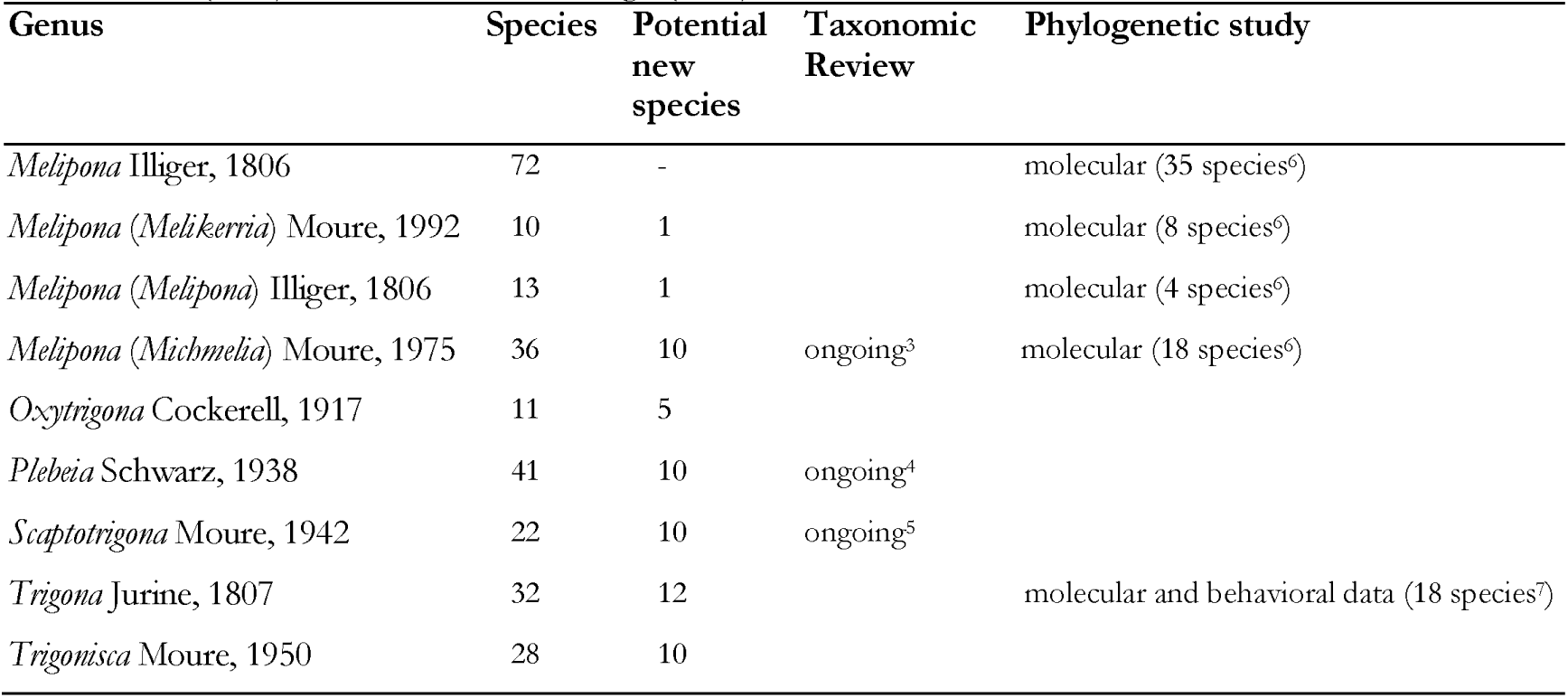
Genera (and subgenera) with ten or more species that demand taxonomic revision, where some have ongoing taxonomic review as indicated. Species number follows 1. Camargo et al. (2023) and potential new species follow Pedro (2014). Ongoing or unpublished taxonomic studies: Melipona (3. JA Santos PhD thesis, INPA); Plebeia (4. H. Werneck & G. Melo) and Scaptotrigona (5. M. Carvalho, master’s Dissertation, FFCLRP) 6. Ramirez et al. (2010); 7. Rasmussen & Camargo (2008).

In addition, some widely distributed species likely represent *species complexes* – groups of closely related taxa with little or no morphological differentiation that are treated under a single name (Table 2). In certain cases, species groups or subspecies have already been recognized, especially for managed species of *Melipona*, such as the Amazonian *Melipona seminigra* (Table 2). However, other commonly managed species—such as *Tetragonisca angustula*, *Cephalotrigona capitata*, and *Scaura longula*—have no formal recognition of species groups or subspecies, despite their large distribution range.

**Table 2.** Species complexes among the managed species of Meliponini in Brazil according to the National Catalogue of Stingless Bees (Portaria N. 665, 3/11/2021). *M. seminigra comprises four subespecies (Camargo et al. 2023). Brazilian states acronyms: AC (Acre), AL (Alagoas), AM (Amazonas), AP (Amapá), BA (Bahia), CE (Ceará), ES (Espírito Santo), GO (Goiás), MA (Maranhão), MG (Minas Gerais), MS (Mato Grosso do Sul), MT (Mato Grosso), PA (Pará), PB (Paraíba), PE (Pernambuco), PI (Piauí), PR (Paraná), RJ (Rio de Janeiro), RO (Rondônia), RR (Roraima), RS (Rio Grande do Sul), SC (Santa Catarina), SE (Sergipe), SP (São Paulo), TO (Tocantins).

| Species names and authors | Geographic distribution (Countries and Brazilian States) |
| --- | --- |
| *Cephalotrigona capitata* (Smith, 1854) | Argentina, Bolivia, Brazil (AP, BA, CE, ES, MT, MG, PR, PA, SC, SP), Colombia, French Guiana, Guiana, Paraguay, Peru, Suriname, Trinidad and Tobago, Venezuela |
| *Melipona* (*Melikerria*) *fasciculata* (Smith, 1854) | Brazil (MA, MT, PA, PI, TO) |
| *Melipona* (*Melikerria*) *interrupta* (Latreille, 1811) | Brazil (AP, AM, PA), French Guiana, Guiana, Suriname |
| *Melipona* (*Michmelia*) *seminigra* Friese, 1903* | Bolivia, Brazil (AC, AM, MA, MT, PA, RO, RR, TO) |
| *Melipona* (*Michmelia*) *fuliginosa* (Lepeletier, 1836) | Argentina, Bolivia, Brazil (AC, AP, AM, BA, ES, GO, MA, MT, PA, PI, RO, RR, SP), French Guiana, Guiana, Suriname, Venezuela |
| *Tetragonisca angustula* (Latreille, 1811) | Mexico, Belize, Bolivia, Brazil (AL, AP, AM, BA, CE, ES, GO, MA, MT, MS, MG, PR, PB, PA, PE, RS, RJ, RR, SC, SE, SP, TO), Colombia, Costa Rica, Ecuador, Guatemala, Guiana, Honduras, Nicaragua, Panama, Peru, Suriname, Venezuela |
| *Plebeia minima* (Gribodo, 1893) | Bolivia, Brazil (AC, AP, AM, MA, MT, PA), Peru, Suriname |
| *Scaura longula* (Lepeletier, 1836) | Bolivia, Brazil (AP, AM, BA, GO, MA, MT, MG, PA, SP, TO), Colombia, Peru, Suriname |

Regarding Darwinian shortfall, 29 genera lack any published phylogenetic study (Table S3), including *Plebeia*, *Trigonisca* or *Scaptotrigona*. Of these, 12 genera contain more than 10 species, most of which are well represented in the Amazon. Most existing phylogenetic studies within Meliponini relied on morphology to infer relationships among species and support taxonomic classification. Some genera, such as *Partamona* and *Geotrigona*, have both phylogenetic analysis and taxonomic revision available. For *Melipona* and *Trigona*, the two most speciose genera, molecular phylogenetic analysis have been conducted. However, due to the taxonomic complexity and richness of these groups, no comprehensive taxonomic treatment currently exists for their full species diversity.

### Geographic distribution and Wallacean shortfalls for managed and non-managed species in the Amazon

Our compiled dataset included 50,548 geographic occurrence records, which reduced to 6,219 unique coordinates after data cleaning (Table S4). These records came from different sources, but 98% correspond to preserved specimens (49,515 records). Most of the data are from collections housed in Brazil and the United States of America (Table S5). Three Brazilian institutions account for 79% of the compiled dataset: RPSP (35,551), DZUP (1,806) and INPA (1,099). The second-largest contributor is the University of Kansas – Snow Entomological Collection (KU SEMC, 5674), which holds approximately 11% of the records, followed by the American Natural History Museum (AMNH, 3,467). All other collections contributed less than 2% each (Table S5).

Figure 2 shows the geographical distribution of Meliponini species across the Amazon biome, including its WWF ecoregions. Some ecoregions are well-represented in terms of recorded occurrence points—for example, Southwestern Amazonian Forests (19,072 occurrences) and Rio Negro-Jurua Moist Forests (11,076 unique coordinates). Others have a moderate number of records (e.g., Tocantins/Pindare moist Forests, 842, and Guianan Moist Forest, 527), while some are poorly sampled (e.g., Xingu-Tocantins-Araguaia Moist Forest, 242, Beni Savanna, 60) (Table S6).

**Figure 2.**
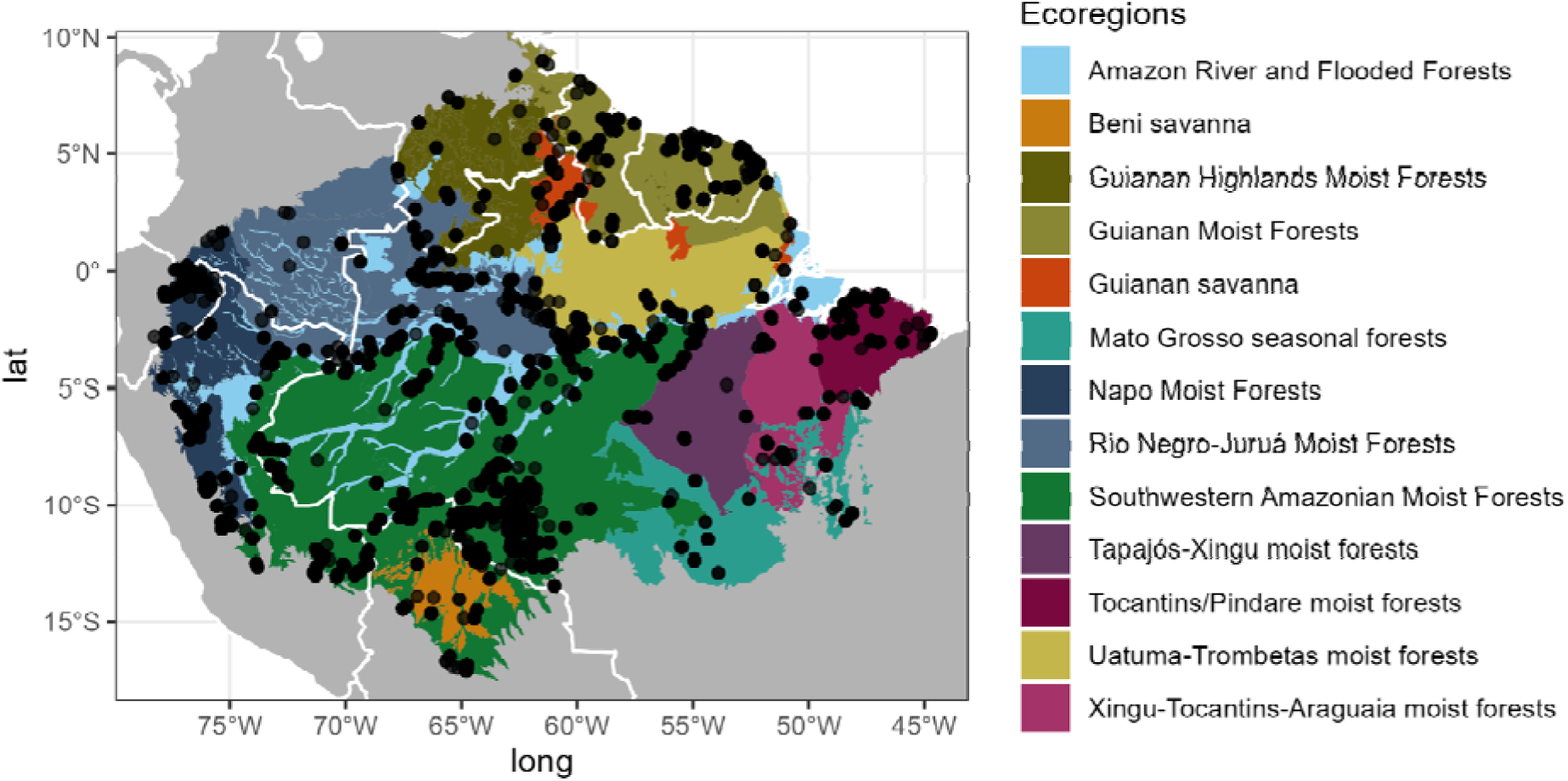
Geographic distribution points for Meliponini in the Amazon Basin over the WWF G200 ecoregions (Olson & Dinerstein 2002). The color legend indicates each of the Amazonian ecoregions.

*Trigona* is the genus with the highest number of unique coordinates (1,940 observations) followed by *Melipona* (1,316 observations), *Partamona* (910 observations) and *Tetragona* (332) (Figure S1). All other regions have fewer than 300 unique records. Some are poorly represented in the databases, such as *Lestrimellita*, *Nogueirapis*, *Leurotrigona*, each with fewer than 50 unique records. Others, such as *Celetrigona*, *Camargoia*, and *Schwarziana*, are represented by fewer than 10 records (Figure S1).

A total of 9,523 records were not identified to species level, corresponding to 1,254 unique coordinates. Among these, *Trigona* (325), *Melipona* (211) and *Tetragona* (196) were the most frequently unidentified genera.

The total number of valid stingless bee species recorded in the geographic databases for the Amazon was 232 species (see *Results, first section*). This number differs by approximately seven species from the total reported for the Amazon in the *Neotropical Bee Catalogue*. This gap suggests that at least seven described species have not been collected again since their original description, have not yet been digitized in the databases we searched, or both.

The complete list of species used in meliponiculture is presented in Table S2 and contains 95 species in 17 genera along with information about its presence in the Amazon. In total, 76 stingless bee species are used in meliponiculture in the Amazon, belonging to 17 genera (Table S2). Similarly to the general distribution pattern of Meliponini in the region, the distribution of managed species is closely associated with major rivers and urbanized areas (Figure 3). Most genera occurring in the Amazon are potentially used for meliponiculture, with the exception of *Oxytrigona*, *Lestrimelitta*, *Nogueirapis*, *Ptilotrigona*, *Trichotrigona* and *Camargoia,* which currently have no known managed species. Some species used in meliponiculture are likely species complexes, and thus require taxonomic revision (Table 2). A well-known example is *Tetragonisca angustula*, commonly known as *jataí*, whose wide geographic distribution suggests it may consist of multiple cryptic species (Table 2). Others, like *Melipona seminigra*, already have formally described subspecies, reinforcing the need for careful taxonomic assessment of managed taxa.

**Figure 3.**
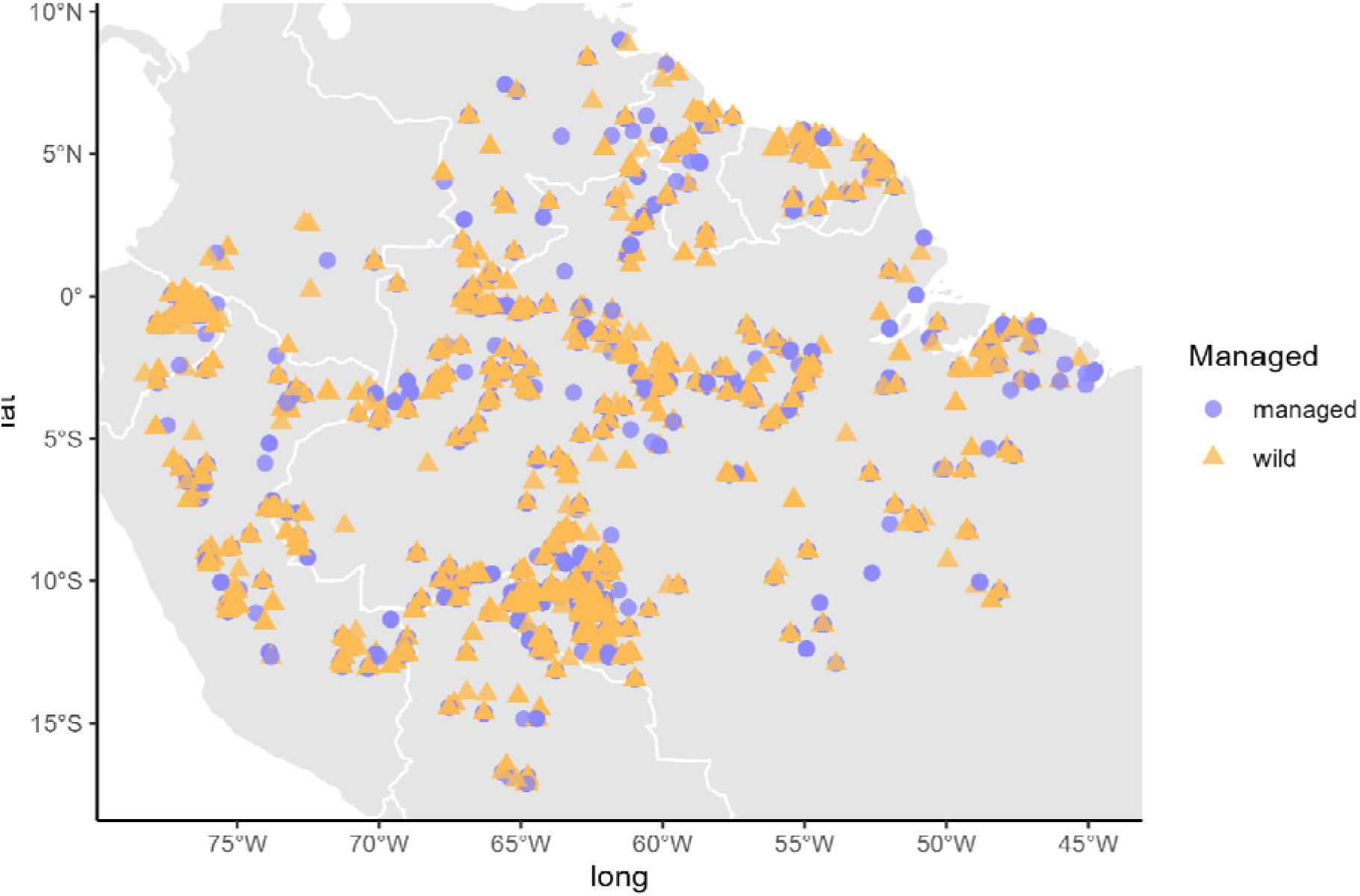
Geographic distribution points for Meliponini highlighting the distribution of managed (yellow triangles) versus non-managed species (purple dots) species.

## Discussion

We identified notable Linnean, Darwinian and Wallacean shortfalls in the knowledge of stingless bees in the Amazon, particularly among the most speciose genera – *Melipona*, *Trigona* – and in widely distributed species complexes such as *Tetragonisca angustula*. Although fewer than a half of the genera and subgenera have been the focus of phylogenetic studies, the relationship among genera are relatively well-known (Rasmussen & Cameron 2010, Lepeco et al. 2024). These studies focused on the entire Meliponini tribe, including lineages from less diverse biogeographic regions such as the Afrotropics and Indo-Australasian realms. Among their main findings are the strong support for a South American origin of stingless bees and the monophyly of lineages found across the three major tropical realms (Rasmussen & Cameron 2010). According to these studies, the Neotropical clade originated in the early Cretaceous (∼80 Ma), with *Trigonisca sensu lato* representing the earliest diverging lineage within the group. Within a broader taxonomic sampling, Lepeco et al. (2024), confirmed the monophyly of most genera and subgenera adopted by recent taxonomic treatments of the Neotropical stingless bees (Camargo et al. 2023, Engel et al. 2023). However, exceptions include *Friseomelitta*, *Plebeia* and *Scaura*, which did not emerge as monophyletic. These results indicate that these three genera, particularly the first two, should be prioritized in future phylogenetic research to support their taxonomic classification. Such efforts are especially urgent given that many undescribed but already recognized species are known to exist within these groups (Pedro 2014).

Among the most taxonomically well-known genera of Meliponini, only a few have been a focus of phylogenetic informed taxonomic studies. Examples include *Geotrigona* Moure, 1943 (Camargo & Moure 1996) and *Partamona* Schwarz, 1939 (Camargo & Pedro 2003). These studies represent the legacy of the late Professor João M. F. Camargo and his collaborators, who adopted a comprehensive approach that integrated morphological, behavioral, and geographical data. As a result, certain genera, such as *Partamona*, have a well-resolved taxonomy supported by a robust set of diagnostic traits, in contrast to other less-studied genera. *Melipona*—the most species-rich genus of stingless bees whose center of diversity is in the Amazon—has been the subject of one major phylogenetic study (Ramírez et al. 2010), which clarified the relationships among its subgenera. These relationships were further supported in the broader tribal-level phylogeny by Lepeco et al. (2024). However, to better understand the origins and diversification of *Melipona*, broader taxonomic sampling is needed. The studies to date have included only a fraction of its diversity (13 species in Lepeco et al. 2024; 35 in Ramírez et al. 2010). *Trigona* is another highly speciose genus that still lacks a complete taxonomic revision, although at least nine species groups have been recognized based on phylogenetic evidence (Rasmussen & Camargo 2008).

Given their importance for meliponiculture and pollination, it is imperative that researchers, conservationists and stingless beekeepers have access to a modern, phylogenetically informed taxonomy that facilitates accurate species recognition. The perspectives are optimistic, as a wide range of source of data is available to support comprehensive and integrative taxonomic studies, like single sequencing, genomic or population genetics data—and have already been successfully applied to vertebrates (e.g., Ribas et al. 2012; Martins et al. 2021), other Hymenoptera (Wang et al. 2016), and other bee groups (Sandoval-Arango et al. 2023; Agostini et al. 2024). While traditional taxonomic and phylogenetic studies have relied heavily on morphological characters of bees and their nests (e.g. Camargo & Pedro 2003), additional data sources have proven highly useful. These include wing morphometrics (e.g. Francoy et al. 2011, Santos et al. 2021), and mitochondrial DNA analysis (Francini et al. 2022, Françoso et al. 2016, 2019). For diverse and widely distributed genera, such as *Melipona* or *Tetragonisca,* an integrative approach —combining morphology, genetic data, and geographic distribution—is essential. This approach not only helps resolve long-standing taxonomic issues, such as the excessive number of synonyms arising from species described without phylogenetic support or based on polymorphic characters, but also enables the robust description of new species supported by multiple lines of evidence.

We identified significant gaps in the knowledge of geographical distribution of stingless bees in the Amazon, which can be attributed to several factors. First, access to existing data from museums and biological collections remains limited. Digitization is one of the bottlenecks contributing to Wallacean shortfalls (Meyer et al. 2015, Marshall et al. 2024), particularly for biodiversity data from tropical regions, which demand substantial research efforts in several areas of knowledge (Diniz-Filho et al. 2023). As observed across many plant and animal groups, a large proportion of tropical ecological research is still conducted by institutions from the Global North, often with limited participation from Global South researchers (Ocampo-Ariza et al. 2023). Moreover, digitizing biological collections is a time– and resource-intensive process that is frequently hindered by a lack of financial or political support. Surprisingly, however, for stingless bees, the majority of our dataset originated from Brazilian institutions, reflecting the efforts of researchers and institutions committed to documenting the country’s biodiversity. Nevertheless, these records represent only a fraction of what the collections actually contain. For example, the RPSP collection houses approximately 150,000 Meliponini specimens, of which about 50% are digitized (<u>SpeciesLink – RPSP</u>). The DZUP collection contains around 350,000 bee specimens, but only about 7% are currently available online (<u>SpeciesLink – DZUP</u>). These figures underscore the urgent need for digitization initiatives in South American biological collections to overcome the Wallacean shortfall in Amazonian Meliponini. Although several governmental and non-governmental programs (e.g., SpeciesLink, SiBBr) have advanced digitization efforts over the past two decades, much work remains to be done.

Secondly, another major source of these gaps is the lack of field collecting in vast regions of the Amazon. Therefore, increased field sampling efforts are essential to address the Wallacean shortfall for stingless bees in this region. Biodiversity knowledge in the Amazon remains heavily influenced by logistical limitations and other human induced factors, such as the absence of research groups in determined areas and complex situations surrounding indigenous territories, which covers about 23% of the Amazon (Carvalho et al. 2023). This bias – driven by accessibility – is a common issue across many other regions and taxa, and has contributed to significant Linnaean, Wallacean, and Hutchinsonian shortfalls (Oliveira et al. 2016). Improving access to underexplored areas could substantially enhance our understanding not only of bees but of Amazonian biodiversity as a whole. Many studies on other animal groups have shown that regions separated by rivers often harbor endemic species, due to the role of rivers as barriers to gene flow (Haffer 1969; Haffer 2008, Guayasamin et al. 2024). In the case of stingless bees, large rivers act as effective barriers to dispersal, as swarming typically occurs no more than 300 meters from the mother colony (Camargo & Pedro 2003; Grüter 2020). This reinforces the importance of conducting fine-scale biogeographic research and local sampling in areas divided by river systems.

In addition to the logistical difficulties of sampling in remote regions of the Amazon, bee sampling poses its own set of challenges. Bee surveys are still largely based on collecting individuals while visiting flowers typically through hand netting, although more comprehensive monitoring is best achieved using a combination of methods (Klaus et al. 2024). Some bee groups are easily sampled using efficient passive techniques, such as the use of trap-nests for cavity-nesting bees (Kamke et al. 2011) or scent baits for attracting orchid bees (Brown et al. 2024). Colored pan traps, a common tool for capturing a wide range of bee taxa in open habitats, have shown limited effectiveness in tropical forest environments, and their utility in such contexts is questioned (Prado et al. 2017). These limitations highlight the need for methodological refinement and context-specific sampling strategies, particularly for effective bee biodiversity surveys in the Amazon.

Sampling bees in the forest canopy remains a major challenge in both temperate and tropical ecosystems. In the Amazon, the canopy represents a vertical frontier of biodiversity research, with enormous potential to uncover high levels of undescribed taxonomic diversity. As emblematic members of tropical bee communities, stingless bees are known to specialize in foraging on tall canopy trees with (Vasconcelos et al. 2018) and tend to prefer mass-flowering trees over herbs or shrubs—even when the latter are more readily accessible (Martins et al. 2023). For this reason, accessing the canopy is critical for effectively sampling Meliponini. Locating nests is also notoriously difficult, given the low nest density and inconspicuous nature of many species’ nests (Oliveira et al. 1995, Roubik 2006). Thus, the development of canopy-specific collecting methods and protocols is essential (e.g. Brown et al. 2024). In other regions, passive traps placed in the canopy (Allen & Davis 2022) and extended entomological nets (Dorey et al. 2024) have been proved successful in sampling canopy bee fauna. When applied in the Amazon, such methods could help reveal bee biodiversity that has been consistently underestimated due to extensive sampling bias (Orr et al. 2021; Roubik 1992). Additionally, chemical attractants— widely used for sampling orchid bees—have also shown promising results for other bee groups (Rabeschini et al. 2021), offering another potential tool for improving canopy sampling.

Lastly, there is a clear overlap between Linnean and Wallacean shortfalls, particularly when occurrence records lack species-level identification. This is especially common in more speciose genera, such as *Trigona*, which limits the utility of these records for species-level analyses. Conversely, there are many cases where taxonomic information is available—the species is described, valid, and sometimes even well-documented in the literature (e.g., Camargo et al. 2023)—but the geographic occurrence data are missing from public databases. Often, these records are known to exist in local museum collections, but they remain undigitized and are therefore absent from large data aggregators. For example, the Lacunas data platform reveals that out of all stingless bee species recorded for Brazil in the *Neotropical Bee Catalogue* (Camargo et al. 2023), 168 species have no occurrence records available in the SpeciesLink network, and 41 species have only 1 to 5 records (Gaps in knowledge of bees in Brazil 2025). This highlights the critical importance of digitizing local collections to bridge both Linnaean and Wallacean knowledge gaps.

Systematic bee surveys and monitoring programs are widely used tools for expanding geographic and taxonomic knowledge and providing essential data for conservation planning (Klaus et al. 2024). However, such efforts remain sporadic in the Amazon, with only a few published studies (e.g., Oliveira et al. 1995; Brown & Albrecht 2001; Silveira et al. 2005; Brown & Oliveira 2014). This limited coverage can be partially attributed to the methodological and logistical challenges of sampling bees in tropical forests, as discussed earlier. Forest fragmentation can have particularly negative impact on some stingless bee species, especially those requiring large population sizes to avoid inbreeding and the resulting production of diploid males, which are eaten by workers, thereby decreasing colony survival (Oliveira et al. 1995). In addition to species-specific behavior, the patchy distribution of stingless bees nests means that some species may respond poorly to habitat disturbance, while others show greater resilience (Brown & Oliveira 2014). Moreover, the low efficiency of traditional sampling methods in forested areas (Oliveira 2001) can bias interpretations of how deforestation affects stingless bee populations. In contrast, orchid bees, whose males are easily attracted with scent baits, have shown marked declines in both species richness and abundance following agricultural settlement in the Amazon (Brown et al 2024). Despite the ecological importance of stingless bees as pollinators, we still lack a comprehensive understanding of how land-use change affects their populations in the Amazon. This remains a critical priority for future research in the region.

Among the species used for meliponiculture, the biodiversity knowledge shortfalls remain pronounced and urgent, especially given the huge potential of these bees to support local development through the production of honey and other products (Quezada-Eúan & Alves 2020, Jaffe et al. 2015). The genera *Melipona*, *Scaptotrigona*, and *Tetragonisca* are the most commonly used in meliponiculture (Quezada-Eúan & Alves 2020), with Melipona standing out in the Amazon Basin—where 24 species are considered suitable, including 21 found exclusively in the state of Amazonas (Menezes et al. 2023). Recognizing these units is fundamental for the development of meliponiculture, which continues to face challenges due to taxonomic uncertainties, lack of standardization, and limited technical knowledge (Jaffé et al., 2015, Carvalho-Zilse & Nunes-Silva 2012). Several species commonly managed in meliponiculture likely represent species complexes—widely distributed taxa composed of multiple, closely related but undescribed species. The best example is the popularly known *jatai*, *Tetragonisca angustula*, which occurs from Mexico to southern Brazil (Camargo et al. 2023). Despite studies on its genetics, behavior and morphology (Cunha et al. 2024, Ferrari et al. 2024), there is still no comprehensive taxonomic revision of this species across its full range. This gap significantly hampers management decisions and limits our understanding of how to optimize meliponiculture in different ecological and cultural contexts.

It is worth noting that the Wallacean shortfall may be less pronounced among managed genera, as those species are often more actively sought after by stingless beekeepers and the target of scientific research. However, the geographic distribution of managed species is frequently biased by the translocation of colonies across regions, including introductions outside their natural distribution range (Jaffe et al. 2015). Despite recent efforts to regulate and standardize the movement of colonies among stingless beekeepers (Brasil 2021), those practices remain common. This widespread translocation introduces important caveats to our interpretation of native distribution patterns and also has negative consequences for meliponiculture (Gonzalez et al. 2022). As with other native species introduced into non-native areas, introduced stingless bees may disrupt local bee communities, compete for resources, or even spread pathogens (Goulson 2003). Therefore, this distributional bias must be taken into account in any assessment of the geographic range of managed stingless bee species, especially when modeling their future distributions under climate and land-use change (Lima & Marchioro 2021).

For example, little is known about how commonly managed stingless bee species will respond to current and future climatic changes, or how those changes might affect their pollination services (Carvalho-Zilse & Nunes-Silva 2012). In the Amazon, there is a significant knowledge gap regarding both the floral resources preferred by these bees and the spatial foraging range required for optimal colony development. Even when colonies are kept near natural areas, they need supplementation with additional floral resources to mitigate the negative effects of intra– and interspecific competition, especially in areas of high colony density (Krug et al. 2021). Notably, even in regions with relatively low bee diversity, stingless bees remain the most common pollinators of flowering plants (Martins et al., in review). Like other eusocial bees, Meliponini maintain perennial colonies, which translates into a high and continuous demand for floral resources, sustaining interactions with a broad array of plant families (Bueno et al. 2023, Ramalho 2004, Martins et al. 2023, Absy et al. 2018). In other tropical biomes, the woody plants dominate the floral resources visited by these bees, especially in the canopy, where Meliponini tend to forage preferentially (Ramalho 2004; Martins et al. 2023). This pattern likely also applies to the Amazon, but the lack of targeted canopy sampling and species-specific floral records hampers deeper conclusions. Because of the difficulty in accessing interactions in the canopy, most of what we know about Meliponini–plant interactions in the Amazon comes from palynological studies of pollen and honey stored in nests (Absy et al. 2018 and references therein). These indirect methods have been crucial in advancing our understanding of plant-bee relationships in the region.

## Conclusion

In this study, we explored the extent of Linnean, Wallacean and Darwinian knowledge gaps for stingless bees in their center of diversity, the Amazon Region. As observed for other insect groups in tropical regions, these gaps are pronounced for stingless bees, limiting our ability to draw conclusions about the effects of global changes on these species (Giannini et al. 2020). Addressing these gaps is increasingly urgent, especially considering the accelerated changes occurring in the Amazon (Esquivel-Muelbert et al. 2019). For stingless bees, these knowledge shortfalls also pose a real barrier for their full integration in sustainable development initiatives. Although they have significant potential to support local economies and livelihoods—particularly for traditional communities that rely on forest-based activities—the lack of basic biological information hampers their effective use. Furthermore, we still know relatively little about how Amazonian forests are pollinated by bees, particularly by stingless bees. Our limited understanding of species diversity and bee-plant interactions across large regions makes it difficult to assess the vulnerability of this essential ecosystem service.

These knowledge gaps stem in part from political and economic factors, which influences both access to existing biological data and the infrastructure required for fieldwork in remote areas. This leads to two major issues: the inaccessibility of data already collected and a lack of new data due to research being concentrated in more accessible or urban regions. On a positive note, recent biodiversity data-sharing initiatives — especially those led by Brazilian institutions — have begun to increase the accessibility of specimen-based data. This may represent a shift away from the historical imbalance in which collections housed in the Global North held most of the information about biodiversity in the Global South, particularly for tropical biomes. We emphasize the urgent need for investment in biological collections, data infrastructure, and research initiatives in tropical regions, with the active participation of scientists from the Global South. Only through this can we meaningfully expand our knowledge and reduce persistent global inequities in biodiversity science. Finally, the impacts of climate change and accelerated land use in the Amazon are likely to affect both wild stingless bee populations and those managed through meliponiculture (Giannini et al. 2017, 2020). In a future where species are better delimited and understood, we will be in a stronger position to investigate ecological interactions, anticipate future shifts, and promote sustainable development strategies that include stingless beekeeping as a conservation-compatible practice.

## Supplementary figures

**Figure S1.**
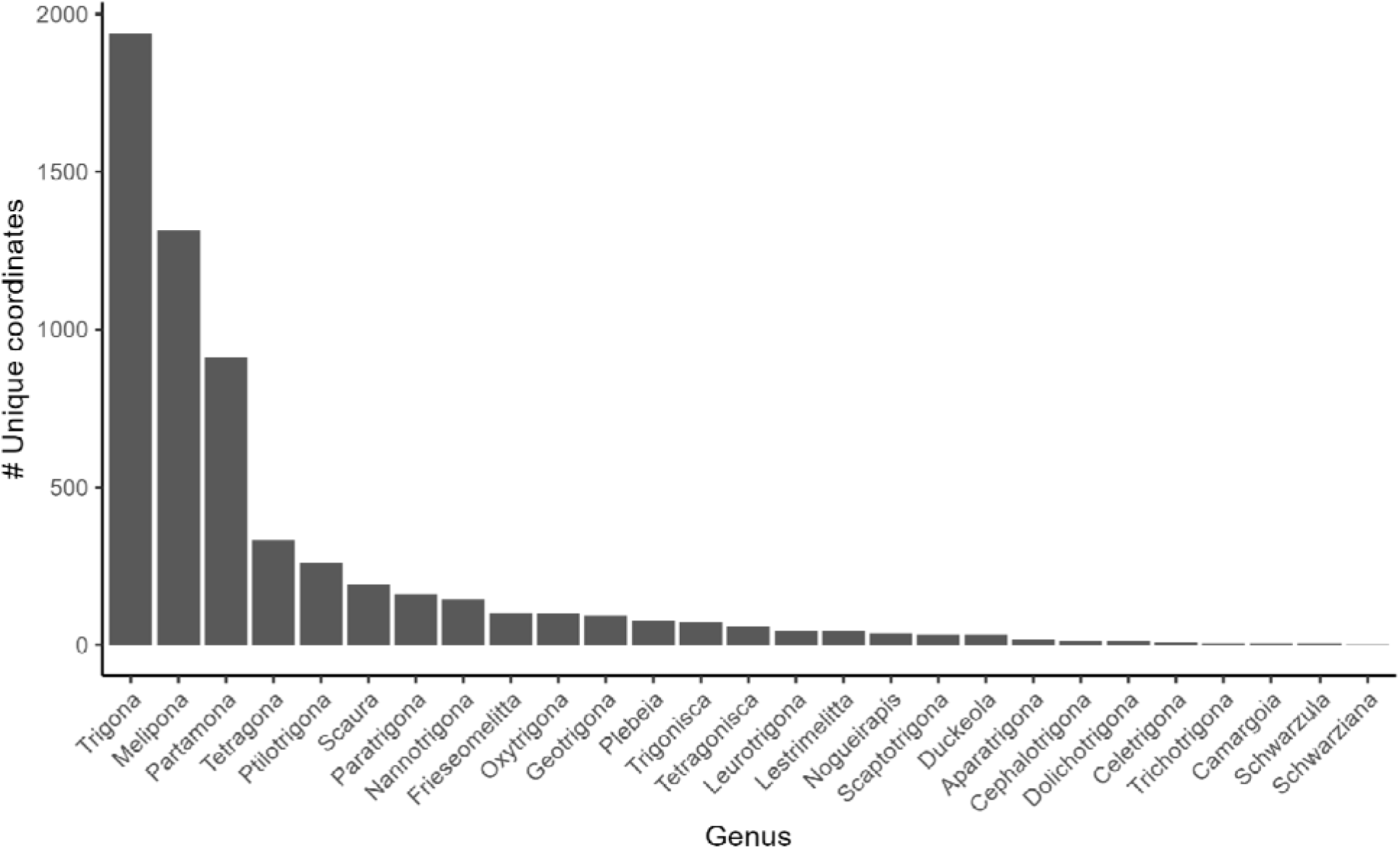
Number of unique coordinates per genus of Meliponini in the Amazon organized from the genus with the highest number of records (Trigona) to the lowest number of records (Schwarziana).

## Notes

### Competing Interest Statement

The authors have declared no competing interest.

## References

1. Absy, M.L., Rech, A.R. & Ferreira, M.G. (2018) Pollen collected by stingless bees: a contribution to understanding Amazonian biodiversity. Pot-Pollen in Stingless Bee Melittology (ed. by P. Vit, S.R.M. Pedro, and D.W. Roubik), pp. 29–46. Springer.

2. Agostini JC, Françoso E, Arias MC, Zanella FCV 2024 Morphological and molecular evidence for considering Xylocopa nigrocincta as the senior synonym of Xylocopa suspecta (Apidae: Xylocopini). Apidol 55:18.

3. Allen G, Davies RG 2023 Canopy sampling reveals hidden potential value of woodland trees for wild bee assemblages. Ins Conserv Div 16:33–46. 10.1111/icad.12606

4. Baranzelli MC, Villalobos F, Cordier JM, Nori J 2023 Knowledge shortfalls’ interactions shadow our perception of species’ exposure to human threats. Biol Conserv 282:110069.

5. Bawa KS, Bullock SH, Perry DR, Coville RE, Grayum MH 1985 Reproductive biology of tropical lowland rain forest trees. II. Pollination systems. Am J Bot 72:346–356.

6. Brasil 2021 Catálogo Nacional de Abelhas-Nativas-Sem-Ferrão. Available at: https://in.gov.br/web/dou/-/portaria-n-665-de-3-de-novembro-de-2021-357715380

7. Brown JC, Corrêa-Neto JDJ, Ribeiro CF, Oliveira ML 2024 The impact of agricultural colonization and deforestation on orchid bees (Apidae: Euglossini) in the Brazilian Amazon. Biol Conserv 293:110560.

8. Bueno FGB, Kendall L, Alves DA, Tamara ML, Heard T, Latty T, Gloag R 2023 Stingless bee floral visitation in the global tropics and subtropics. Global Ecol Conserv 43:e02454.

9. Camargo JMF, Pedro SRM, Melo GAR 2023 Meliponini Lepeletier, 1836. Catalogue of Bees (Hymenoptera, Apoidea) in the Neotropical Region – online version.

10. Camargo JMF, Pedro SRM 2003 Meliponini neotropicais: o gênero Partamona Schwarz, 1939 (Hymenoptera, Apidae, Apinae) – bionomia e biogeografia. Revista Brasileira de Entomologia 47:311–372.

11. Camargo JMF, Moure JS 1996 Meliponini neotropicais: o gênero Geotrigona Moure, 1943 (Apinae, Apidae, Hymenoptera), com especial referência à filogenia e biogeografia. Arq Zoo Univ de São Paulo 33:95–161.

12. Cardoso P, Erwin TL, Borges PAV, New TR 2011 The seven impediments in invertebrate conservation and how to overcome them. Biol Cons 144:2647–2655.

13. Carvalho, R.L… & Ferreira, J. (2023) Pervasive gaps in Amazonian ecological research. Current Biology 33:3495–3504.e4.

14. Carvalho-Zilse GA, Nunes-Silva CG 2012 Threats to the stingless bees in the Brazilian Amazon: how to deal with scarce biological data and an increasing rate of destruction. In: Florio RM (ed.) Bees: biology, threats and colonies. Nova Science Publishers.

15. Cunha M, Moura Novaes C, Amorim Pereira J, Mapingala Capoco M, Fernandes-Salomão TM, Meneses Lopes D 2023 Supernumerary B Chromosomes of Tetragonisca fiebrigi share repeat content with standard chromosome set of both T. fiebrigi and Tetragonisca angustula (Apidae: Meliponini). Cytogenetic and Genome Research 163:52–58.

16. Chamberlain S, Barve V, Mcglinn D, Oldoni D, Desmet P, Geffert L, Ram K 2025 rgbif: Interface to the Global Biodiversity Information Facility API. R package version 3.8.1.2. https://CRAN.R-project.org/package=rgbif

17. CRIA – Centro de Referência em Informação Ambiental. (n.d.) speciesLink: Distributed Information System for Biological Collections. Retrieved June 2024, from http://splink.cria.org.br/index?&setlang=en

18. Diniz-Filho JAF, Jardim L, Guedes JJM, Meyer L, Stropp J, Frateles LEF, Pinto RB, Lohmann LG, Tessarolo G, de Carvalho CJB, Ladle RJ, Hortal J 2023 Macroecological links between the Linnean, Wallacean, and Darwinian shortfalls. Front Biogeog 15(2):e59566

19. Dorey JB et al 2024 Canopy specialist Hylaeus bees highlight sampling biases and resolve Michener’s mystery. Front Ecol Evol 12. 10.3389/fevo.2024.1339446

20. Dinerstein E, Olson D, Joshi A, Vynne C, Burgess ND, Wikramanayake E, et al. (2017) An ecoregion-based approach to protecting half the terrestrial realm. BioScience 67(6):534–545.

21. Engel MS, Rasmussen C, Ayala R, De Oliveira FF (2023) Stingless bee classification and biology (Hymenoptera, Apidae): a review, with an updated key to genera and subgenera. ZooKeys 1172:239–312.

22. Entomological Society of America (2021) ESA Position Statement on Insects and Biodiversity. Accessed on February 5th, 2025.

23. Esquivel-Muelbert A, et al. (2019) Compositional response of Amazon forests to climate change. Global Change Biology 25:39–56.

24. Ferrari RR, Ricardo PC, Dias FC, de Souza Araujo N, Soares DO, Zhou Q-S, Zhu C-D, Coutinho LL, Arias MC, Batista TM (2024) The nuclear and mitochondrial genome assemblies of Tetragonisca angustula (Apidae: Meliponini), a tiny yet remarkable pollinator in the Neotropics. BMC Genomics 25:587.

25. Francoy TM, et al. (2011) Geometric morphometrics of the wing as a tool for assigning genetic lineages and geographic origin to Melipona beecheii (Hymenoptera: Meliponini). Apidol 42:499–507. 10.1007/s13592-011-0013-0

26. Francini IB, et al. (2022) DNA barcoding reveals long-term speciation processes in subspecies of the Melipona (Michmelia) seminigra complex (Hymenoptera: Apidae). Eur J Ent 119:309–317. 10.14411/eje.2022.032

27. Françoso E, et al. (2016) A protocol for isolating insect mitochondrial genomes: a case study of NUMT in Melipona flavolineata (Hymenoptera: Apidae). Mit DNA Part A 27:2401–2404. 10.3109/19401736.2015.1028049

28. Françoso E, et al. (2019) Conserved numts mask a highly divergent mitochondrial-COI gene in a species complex of Australian stingless bees Tetragonula (Hymenoptera: Apidae). Mit DNA Part A 30:806–817. 10.1080/24701394.2019.1665036

29. Gaps in Knowledge of Bees in Brazil. Available at: https://moure.cria.org.br/lacunas/202501 (Accessed 15 July 2025).

30. Grüter C (2020) Stingless Bees: Their Behaviour, Ecology and Evolution. Springer International Publishing, Cham.

31. Guayasamin JM, Ribas CC, Carnaval AC, Carrillo JD, Hoorn C, Lohmann LG, Riff D, Ulloa Ulloa C, Albert JS (2024) Evolution of Amazonian biodiversity: A review. Acta Amazonica 54:e54bc21360.

32. Giannini TC, Costa WF, Borges RC, Miranda L, da Costa CPW, Saraiva AM, Imperatriz Fonseca VL (2020) Climate change in the Eastern Amazon: crop-pollinator and occurrence-restricted bees are potentially more affected. Regional Environmental Change 20:9.

33. Giannini TC, Costa WF, Cordeiro GD, Imperatriz-Fonseca VL, Saraiva AM, Biesmeijer J, Garibaldi LA (2017) Projected climate change threatens pollinators and crop production in Brazil. PLOS ONE 12:e0182274.

34. Gonzalez VH, Oyen K, Vitale N, Ospina R (2022) Neotropical stingless bees display a strong response in cold tolerance with changes in elevation. Conservation Physiology 10:coac073.

35. Goulson D (2003) Effects of introduced bees on native ecosystems. Annu Rev Ecol Evol Syst 34:1–26.

36. Haffer J (2008) Hypotheses to explain the origin of species in Amazonia. Braz J Biol 68:917–947.

37. Haffer J (1969) Speciation in Amazonian forest birds. Science 165:131–137.

38. Hortal J, De Bello F, Diniz-Filho JAF, Lewinsohn TM, Lobo JM, Ladle RJ (2015) Seven shortfalls that beset large-scale knowledge of biodiversity. Annu Rev Ecol Evol Syst 46:523–549.

39. Instituto Brasileiro de Geografia e Estatística (IBGE) (2019) Biomas do Brasil – Visão geral. Available at: https://www.ibge.gov.br/apps/biomas/#/home (Accessed July 10 2025).

40. Kamke R, Zillikens A, Steiner J (2011) Species richness and seasonality of bees (Hymenoptera, Apoidea) in a restinga area in Santa Catarina, southern Brazil. Stud Neotrop Fauna Environ 46:35–48.

41. Klaus F, Ayasse M, Classen A, Dauber J, Diekötter T, Everaars J, Fornoff F, Greil H, Hendriksma HP, Jütte T, Klein AM, Krahner A, Leonhardt SD, Lüken DJ, Paxton RJ, Schmid-Egger C, Steffan-Dewenter I, Thiele J, Tscharntke T, Erler S, Pistorius J (2024) Improving wild bee monitoring, sampling methods, and conservation. Basic Appl Ecol 75:2–11.

42. Krug C, Gomes FB, Oliveira MM, Ferreira GAC (2021) Plantas para a meliponicultura na Amazônia: guia ilustrado. Embrapa Amazônia Ocidental, Brasília.

43. Lacunas de conhecimento das Abelhas do Brasil. Available at: https://moure.cria.org.br/lacunas/202402 (Accessed 1 July 2025).

44. Lepeco A, Branstetter MG, Melo GAR, Freitas FV, Tobin KB, Gan J, Jensen J, Almeida EAB (2024) Phylogenomic insights into the worldwide evolutionary relationships of the stingless bees (Apidae, Meliponini). Mol Phyl Evol 201:108219.

45. Jaffé R, Pope N, Carvalho AT, Maia UM, Blochtein B, De Carvalho CAL, Carvalho-Zilse GA, Freitas BM, Menezes C, De Fátima Ribeiro M, Venturieri GC, Imperatriz-Fonseca VL (2015) Bees for development: Brazilian survey reveals how to optimize stingless beekeeping. PLOS ONE 10:e0121157.

46. Marshall L, Leclercq N, Carvalheiro LG, Dathe HH, Jacobi B, Kuhlmann M, Potts SG, Rasmont P, Roberts SPM, Vereecken NJ (2024) Understanding and addressing shortfalls in European wild bee data. Biol Conserv 290:110455.

47. Martins AC, Melo GAR, Renner SS (2014) The corbiculate bees arose from New World oil-collecting bees: implications for the origin of pollen baskets. Mol Phyl Evol 80:88–94.

48. Martins AC, Heinrich L, Hughes AC, Seltmann KC, Orr MC, Vasconcelos T (in review) Plant communities in the Americas are highly bee dependent regardless of biome or local bee diversity.

49. Martins AC, et al. (2023) Contrasting patterns of foraging behavior in neotropical stingless bees using pollen and honey metabarcoding. Sci Rep 13:14474. 10.1038/s41598-023-41304-0

50. Martins LF, et al. (2021) Whiptail lizard lineage delimitation and population expansion as windows into the history of Amazonian open ecosystems. Syst Biodiv 19:957–975. 10.1080/14772000.2021.1953185

51. Melo GAR (2020) Stingless Bees (Meliponini). In: Starr CK (ed) Encyclopedia of Social Insects, pp. 1–18. Springer International Publishing, Cham.

52. Menezes C, Alves DA, Lucena DAA, Almeida EAB (2023) Abelhas sem ferrão relevantes para a meliponicultura no Brasil. ABELHA, São Paulo, SP.

53. Michener CD (2007) The Bees of the World, 2nd ed. Johns Hopkins University Press, Baltimore.

54. Ocampo-Ariza C, Toledo-Hernández M, Librán-Embid F, Armenteras D, Vansynghel J, Raveloaritiana E, Arimond I, Angulo-Rubiano A, Tscharntke T, Ramírez-Castañeda V, Wurz A, Marcacci G, Anders M, Urbina-Cardona JN, de Vos A, Devy S, Westphal C, Toomey A, Sheherazade, Chirango Y, Maas B (2023) Global South leadership towards inclusive tropical ecology and conservation. Perspect Ecol Conserv 21:17–24.

55. Olson DM, Dinerstein E, Wikramanayake ED, Burgess ND, Powell GVN, Underwood EC, D’Amico JA, Itoua I, Strand HE, Morrison JC, Loucks CJ, Allnutt TF, Ricketts TH, Kura Y, Lamoreux JF, Wettengel WW, Hedao P, Kassem KR (2001) Terrestrial ecoregions of the world: a new map of life on Earth. BioScience 51:933–938.

56. Oliveira U, Paglia AP, Brescovit AD, De Carvalho CJB, Silva DP, Rezende DT, Leite FSF, Batista JAN, Barbosa JPPP, Stehmann JR, Ascher JS, De Vasconcelos MF, De Marco P, LöwenbergCNeto P, Dias PG, Ferro VG, Santos AJ (2016) The strong influence of collection bias on biodiversity knowledge shortfalls of Brazilian terrestrial biodiversity. Divers Distrib 22:1232–1244.

57. Oliveira ML (2001) Stingless bees (Meliponini) and orchid bees (Euglossini) in “terra firme” tropical forests and forest fragments. In: Bierregaard RO Jr, Gascon C, Lovejoy TE, Mesquita R (eds) Lessons from Amazonia: The Ecology and Conservation of a Fragmented Forest, pp. 208–218. Yale University Press, New Haven.

58. Oliveira ML, Morato EF, Garcia MVB (1995) Diversidade de espécies e densidade de ninhos de abelhas sociais sem ferrão (Hymenoptera, Apidae, Meliponinae) em floresta de terra firme na Amazônia central. Rev Bras Zool 12:13–24.

59. Olson DM, Dinerstein E (2002) The Global 200: priority ecoregions for global conservation. Ann Mo Bot Gard 89(2):199–224.

60. Ollerton J, Winfree R, Tarrant S (2011) How many flowering plants are pollinated by animals? Oikos 120:321–326.

61. Orr MC, Hughes AC, Chesters D, Pickering J, Zhu C-D, Ascher JS (2021) Global patterns and drivers of bee distribution. Curr Biol 31:451–458.e4.

62. Pebesma E, Bivand R (2023) Spatial Data Science: With Applications in R. Chapman and Hall/CRC. 10.1201/9780429459016

63. Pebesma E (2018) Simple Features for R: standardized support for spatial vector data. R J 10(1):439–446. 10.32614/RJ-2018-009

64. Pedro SRM (2014) The stingless bee fauna in Brazil (Hymenoptera: Apidae). Sociobiology 61:348– 354.

65. Prado SG, Ngo HT, Florez JA, Collazo JA (2017) Sampling bees in tropical forests and agroecosystems: a review. J Insect Conserv 21:753–770.

66. Quezada-Euán JJG, Alves DA (2020) Meliponiculture. In: Starr CK (ed) Encyclopedia of Social Insects, pp. 587–592.

67. R Core Team (2021) R: A Language and Environment for Statistical Computing. R Foundation for Statistical Computing, Vienna, Austria.

68. Ramalho M (2004) Stingless bees and mass flowering trees in the canopy of Atlantic Forest: a tight relationship. Acta Bot Bras 18:37–47.

69. Rabeschini G, Joaquim Bergamo P, Nunes CEP (2021) Meaningful words in crowd noise: searching for volatiles relevant to carpenter bees among the diverse scent blends of bee flowers. J Chem Ecol 47:444–454.

70. Ramírez SR, Nieh JC, Quental TB, Roubik DW, Imperatriz-Fonseca VL, Pierce NE (2010) A molecular phylogeny of the stingless bee genus Melipona (Hymenoptera: Apidae). Mol Phyl Evol 56:519–525.

71. Rasmussen C, Camargo JMF (2008) A molecular phylogeny and the evolution of nest architecture and behavior in Trigona s.s. (Hymenoptera: Apidae: Meliponini). Apidologie 39:102–118.

72. Rasmussen C, Cameron SA (2010) Global stingless bee phylogeny supports ancient divergence, vicariance, and long distance dispersal. Biol J Linn Soc 99:206–232.

73. Ribas CC, et al. (2011) A palaeobiogeographic model for biotic diversification within Amazonia over the past three million years. Proc R Soc B 279:681–689. 10.1098/rspb.2011.1120

74. Roubik DW (2006) Stingless bee nesting biology. Apidologie 37:124–143.

75. Roubik DW (2023) Stingless bee (Apidae: Apinae: Meliponini) ecology. Annu Rev Entomol 68:231–256.

76. Roubik DW (1992) Loose niches in tropical communities: why are there so few bees and so many trees? In: Hunter M (ed) Effects of Resource Distribution on Animal–Plant Interactions, pp. 327–354. Elsevier. 10.1016/B978-0-08-091881-5.50014-1

77. R Core Team (2024) R: A Language and Environment for Statistical Computing. R Foundation for Statistical Computing, Vienna, Austria. https://www.R-project.org/

78. Sandoval-Arango S, et al. (2023) Phylogenomics reveals within-species diversification but incongruence with color phenotypes in widespread orchid bees (Hymenoptera: Apidae: Euglossini). Insect Syst Divers 7:1. 10.1093/isd/ixad005

79. Santos IS, Nogueira DS, Castro I, Teixeira JSG, Freitas GS, Oliveira ML (2021) Padrões morfológicos na venação alar de espécies de Tetragona Lepeletier & Serville, 1828 do grupo clavipes (Hymenoptera: Apidae: Meliponini). Entomol Commun 3:ec03032.

80. Silveira OT, Esposito MC, Santos JND, Gemaque FE (2005) Social wasps and bees captured in carrion traps in a rainforest in Brazil. Entomol Sci 8:33–39.

81. SiBBR – Sistema de Informação sobre a Biodiversidade Brasileira. Available at: https://www.sibbr.gov.br (Accessed July 2025).

82. Souza C, et al. (2020) Reconstructing three decades of land use and land cover changes in Brazilian biomes with Landsat archive and Earth Engine. Remote Sens 12(17):2735. 10.3390/rs12172735

83. van Dulmen A (2001) Pollination and phenology of flowers in the canopy of two contrasting rain forest types in Amazonia, Colombia. Plant Ecol 153:73–85.

84. Vasconcelos TNC, Chartier M, Prenner G, Martins AC, Schönenberger J, Wingler A, Lucas E (2019) Floral uniformity through evolutionary time in a speciesCrich tree lineage. New Phytol 221:1597–1608.

85. Wang Y, Zhou Q-S, Qiao H-J, Zhang A-B, Yu F, Wang X-B, Zhu C-D, Zhang Y-Z (2016) Formal nomenclature and description of cryptic species of the Encyrtus sasakii complex (Hymenoptera: Encyrtidae). Sci Rep 6:34372. 10.1038/srep34372

86. Wickham H, Averick M, Bryan J, Chang W, McGowan L, François R, Grolemund G, Hayes A, Henry L, Hester J, Kuhn M, Pedersen T, Miller E, Bache S, Müller K, Ooms J, Robinson D, Seidel D, Spinu V, Yutani H (2019) Welcome to the tidyverse. J Open Source Softw 4(43):1686. 10.21105/joss.01686

87. Zizka A, Silvestro D, Andermann T, Azevedo J, Duarte Ritter C, Edler D, Farooq H, Herdean A, Ariza M, Scharn R, Svanteson S, Wengtrom N, Zizka A, Antonelli A (2019)

88. CoordinateCleaner: standardized cleaning of occurrence records from biological collection databases. Methods Ecol Evol 10(5):744–751. 10.1111/2041-210X.13152

